# The age-dependent relationship between resting heart rate variability and functional brain connectivity

**DOI:** 10.1101/360909

**Authors:** D. Kumral, H.L. Schaare, F. Beyer, J. Reinelt, M. Uhlig, F. Liem, L. Lampe, A. Babayan, A. Reiter, M. Erbey, J. Roebbig, M. Loeffler, M.L. Schroeter, D. Husser, A.V. Witte, A. Villringer, M. Gaebler

## Abstract

Resting heart rate variability (HRV), an index of parasympathetic cardioregulation and an individual trait marker related to mental and physical health, decreases with age. Previous studies have associated resting HRV with structural and functional properties of the brain – mainly in cortical midline and limbic structures. We hypothesized that HRV may alter its relationship with brain structure and function across the adult lifespan. In 388 healthy subjects of three age groups (140 younger: 26.0±4.2 years, 119 middle-aged: 46.3±6.2 years, 129 older: 66.9±4.7 years), gray matter structure (voxel-based morphometry) and resting-state functional connectivity (eigenvector centrality mapping and exploratory seed-based functional connectivity) were related to resting HRV, measured as the root mean square of successive differences (RMSSD). Confirming previous findings, resting HRV decreased with age. For HRV-related gray matter volume, there were no statistically significant differences between the age groups, nor similarities across all age groups. In whole-brain functional connectivity analyses, we found an age-dependent association between resting HRV and eigenvector centrality in the bilateral ventromedial prefrontal cortex (vmPFC), driven by the younger adults. Across all age groups, HRV was positively correlated with network centrality in bilateral posterior cingulate cortex. Seed-based functional connectivity analysis using the vmPFC cluster revealed an HRV-related cortico-cerebellar network in younger but not in middle-aged or older adults. Our results indicate that the decrease of HRV with age is accompanied by changes in functional connectivity along the cortical midline. This extends our knowledge of brain-body interactions and their changes over the lifespan.

## 1. Introduction

Behavioral and physiological changes that occur with advancing age become manifest in the structure and function of multiple macro- and micro-systems of the human organism (Arking, 2006). Important alterations occur in the cardiovascular and the nervous systems, which are coupled to react dynamically to environmental demands (McEwen, 2003). Such adaptations to internal and external challenges, while leaving an imprint on body and brain, underlie healthy aging (Lipsitz and Goldberger, 1992; Swank, 1996). They are also reflected in brain-heart interactions – particularly in parasympathetic cardioregulation – that can be measured as resting heart rate variability (HRV)

HRV quantifies variations in the cardiac beat-to-beat (or RR) interval that can be measured with an electrocardiogram (ECG). Phasic modulation of the heart rate arises from the influences of both branches of the autonomic nervous system (ANS), the parasympathetic (PNS) and the sympathetic nervous system (SNS) – with the PNS quickly lowering the heart rate and the SNS slowly increasing it. Because the PNS has very short response latencies, HRV – and some HRV measures more than others – represents parasympathetic (i.e., vagal) influences on the heart (Thayer and Lane, 2007). HRV, typically acquired at rest, is known to decrease with age (De Meersman and Stein, 2007; Umetani et al., 1998). Preservation of autonomic function, as indexed by relatively increased HRV, has been shown a prerequisite for longevity and healthy aging (Zulfiqar et al., 2010). Higher HRV has also been associated with higher health (Kemp and Quintana, 2013) – for example, with less cardiovascular diseases (Liao et al., 1997; Thayer et al., 2010) – and reduced overall mortality (Buccelletti et al., 2009). In older adults, HRV can indicate inter-individual differences in cognitive performance (Mahinrad et al., 2016; Zeki Al Hazzouri et al., 2014). Hence, resting HRV could be regarded as a biomarker of healthy aging.

The neurovisceral integration model considers the role of the brain in parasympathetic cardioregulation and provides a framework to explain individual differences in resting vagal function (Kemp et al., 2017; Thayer et al., 2012). According to this model, frontal and midbrain areas interact and the prefrontal cortex (PFC) inhibits subcortical regions as well as the ANS. Thereby, the heart is under tonic inhibitory control by the ANS. Assuming this close interaction of the brain and the ANS in heart rate regulation, it has been suggested that inter-individual differences in HRV may reflect structural and functional variability in the brain (Thayer et al., 2012). Indeed, inter-individual differences in resting HRV have been associated with cortical thickness in right anterior midcingulate cortex (aMCC) (Winkelmann et al., 2017) as well as in rostral anterior cingulate cortex (ACC) and left lateral orbitofrontal cortex (OFC) (Yoo et al., 2017). A recent study in individuals between 20 and 60 years found a negative correlation between resting HRV and gray matter volume in limbic structures like insula, amygdala, and parahippocampal gyrus (Wei et al., 2018). Similar brain regions have also been related to HRV in functional neuroimaging studies (Holzman and Bridgett, 2017; Mather and Thayer, 2018; Thayer et al., 2012); both task-based (e.g., BOLD: Critchley et al., 2000; regional cerebral blood flow; rCBF: Gianaros et al., 2004; meta-analyses: Beissner et al., 2013; Thayer et al., 2012) and under resting state conditions (Chang et al., 2013; Jennings et al., 2016; Sakaki et al., 2016). In these studies, activation and connectivity in the medial prefrontal cortex (mPFC), anterior cingulate cortex (ACC), and posterior cingulate cortex (PCC) have most consistently been associated with HRV. These brain areas involved in parasympathetic cardioregulation overlap with the default mode network (DMN) and particularly its nodes along the cortical midline (Beissner et al., 2013). However, the only fMRI study that investigated heart-brain interactions across the adult lifespan included 17 younger and 18 older subjects and restricted their analyses to *a priori* defined regions-of-interest (Sakaki et al., 2016). Across all subjects, higher HRV was related to stronger functional connectivity between right amygdala and medial prefrontal regions, while age group differences were found in HRV-related connectivity between right amygdala and lateral prefrontal regions.

We here investigated brain-heart interactions across the adult lifespan by combining measures of brain structure and function with the assessment of resting HRV. The main aims of this study were to examine (i) the relationship between resting HRV, brain structure, and functional connectivity as well as (ii) its dependence on age in a large sample of healthy adults across the lifespan. Based on previous findings (reviewed above), we hypothesized that inter-individual differences in the brain correlate with inter-individual differences in resting HRV and that this correlation changes with age. To detect HRV-related structural alterations, we used voxel-based morphometry (VBM) (Ashburner and Friston, 2000). To assess HRV-related changes in the functional architecture across the whole brain, we used the graph-based method of eigenvector centrality mapping (ECM). ECM can identify important network nodes (in this case: voxels) based on their functional connectivity (similar to Google’s page rank algorithm) and without the need of an *a priori* selection of a specific seed region or the number of networks / components (Lohmann et al., 2010; Wink et al., 2012). To further explore ECM-derived whole-brain connectivity patterns, we also implemented a resting-state seed-based connectivity analysis (for more details see *Methods*).

## 2. Methods

### 2.1. Participants

Data from two studies were used: (I) the Leipzig Research Centre for Civilization Diseases (LIFE; Loeffler et al., 2015) and (II) the “Leipzig Study for Mind-Body-Emotion Interactions” (LEMON; Babayan et al., under review).

LIFE is a large population-based cohort study from Leipzig, Germany (Loeffler et al., 2015). From the sample of LIFE subjects with MRI data (n = 2,667), we selected healthy subjects between the ages of 20 and 80 years. We applied strict exclusion criteria in three categories: I) health-related criteria; participants were excluded if they reported any medication intake except vitamin food supplements, any past or present cardiovascular health problems and diagnoses, or surgeries, any other medical history and/or diagnosis, in a medical interview. II) ECG-related criteria (see details on ECG acquisition below); if a subject had more than one ECG recording, we used the first acquired ECG file that was collected on the same day as the MRI acquisition. Otherwise, we selected the ECG recording that was temporally closest to MRI acquisition. Regarding data quality, we excluded data with unrepairable signal artifacts or problems regarding R-peak detection. We also omitted data with any abnormal ECG signal (e.g., supraventricular extrasystoles) after visual inspection as well as subjects with extreme HRV values based on Tukey’s (1977) criterion of 3 interquartile ranges (IQR) above the LIFE sample median (N=14, *Median*: 30.13, IQR: 29.76). III) MRI-related criteria; we excluded subjects with incidental findings (e.g., brain tumor, multiple sclerosis, or stroke) on T1-weighted and/or fluid-attenuated inversion recovery (FLAIR) images. We further excluded subjects based on rs-fMRI quality assessment, for example with faulty preprocessing (e.g., during denoising) or excessive head motion (criterion: mean framewise displacement (FD) > 0.6 mm; Power et al., 2012).

LEMON is a cross-sectional sample of healthy younger and older subjects from Leipzig, Germany, who had never participated in another “psychological or MRI research”-related study, did not report any neurological disorders, head injury, any medication affecting the cardiovascular and/or central nervous system, alcohol or other substance abuse, hypertension, pregnancy, claustrophobia, chemotherapy and malignant diseases, current and/or previous psychiatric disease (Babayan et al., under review). The LEMON sample comprised 171 eligible subjects divided into two age groups (young: 20-35 years, old: 59-75 years). Similar to the exclusion criteria mentioned above, subjects with incomplete data (N = 38), incidental findings in MRI (FLAIR, T2-weighted, T1-weighted, SWI) (N=7), or psychoactive drug intake (e.g., tetrahydrocannabinol) determined by urine test (N = 9) were excluded. Two subjects were discarded due to the HRV outlier criterion mentioned above (LEMON sample *Median*: 40.62, IQR: 39.82) and five subjects due to excessive head motion (mean FD > 0.6 mm; Power et al., 2012). To increase the statistical power and the comparability, we pooled the two samples and divided them into three age groups: young (20-35 years from LIFE and LEMON), middle-aged (35-60 years from LIFE), and old (60-80 years from LIFE and LEMON). Details are provided in Table 1.

**Table 1.** Participant characteristics for each age group. For continuous variables, data is provided in means and standard deviations (in parenthesis). One-way ANOVAs were used to detect age group differences.

| | Young<br>(20-35 years)<br>(N=140) | Middle<br>(35-60 years)<br>(N=119) | Old<br>(60-80 years)<br>(N=129) | df | F-value | Eta-<br>squared<br>($\eta^2$) |
| --- | --- | --- | --- | --- | --- | --- |
| Age | 26.01<br>(4.17) | 46.39<br>(6.25) | 66.88<br>(4.68) |  |  |  |
| Sex | 38 M / 102 F | 36 M / 83 F | 50 M / 79 F |  | 4.38‡ |  |
| Resting HRV<br>(RMSSD in ms) | 53<br>(27.11) | 32.77<br>(21.01) | 27.27<br>(22.99) | 385 | 43.01*** | 0.182 |
| Mean HR<br>(1/min) | 64.38<br>(9.62) | 62.93<br>(10.01) | 66.24<br>(10.44) | 385 | 3.411* | 0.017 |
| RR interval (ms) | 952.56<br>(137.06) | 977.87<br>(149.75) | 928.32<br>(148.53) | 385 | 3.62* | 0.018 |
| mean FD (mm) | 0.18<br>(0.05) | 0.28<br>(0.10) | 0.31<br>(0.11) | 385 | 82.61*** | 0.300 |
| BMI (kg/m<sup>2</sup>) | 23.58<br>(3.03) | 26.51<br>(3.62) | 26.54<br>(3.57) | 382 | 33.34*** | 0.148 |
| WHR | 0.86<br>(0.07) | 0.92<br>(0.08) | 0.95<br>(0.08) | 381 | 47.64*** | 0.200 |
| SBP (mmHg) | 122.08<br>(11.42) | 126.55<br>(13.74) | 138.76<br>(18.2) | 381 | 45.45*** | 0.192 |
| DBP (mmHg) | 71.23<br>(7.33) | 78.21<br>(9.11) | 80.00<br>(10.44) | 381 | 35.47*** | 0.156 |
| TMT A (s) | 24.95<br>(7.79) | 30.33<br>(12.73) | 40.01<br>(13.54) | 384 | 58.48*** | 0.233 |
| TMT B (s) | 57.72<br>(17.89) | 71.40<br>(39.04) | 95.15<br>(45.09) | 382 | 45.73*** | 0.193 |

\* $p < 0.05$ ; \*\* $p < 0.01$ ; \*\*\* $p < 0.001$ , 2-tailed
‡ Kruskal-Wallis-Test
*Note.* HRV = heart rate variability; RMSSD = root mean square of successive differences; HR = heart rate, FD = framewise displacement; BMI = body mass index; WHR = waist to hip ratio; SBP = systolic blood pressure; DBP = diastolic blood pressure; TMT = trail making test.

Both studies were in agreement with the Declaration of Helsinki and approved by the ethics committee of the medical faculty at the University of Leipzig, Germany.

### 2.2. ECG collection and HRV analysis

#### LIFE sample

Ten seconds of a standard medical 12-lead resting ECG were acquired using a Page-Writer TC50 ECG system (Philips Medical Systems, Amsterdam, Netherlands) in supine position. We used lead I (from Einthoven’s triangle) for the analysis. R-peaks were automatically detected using the findpeaks function in Matlab 9 (The MathWorks, Inc., Natick, Massachusetts) or Kubios 2.2 (Tarvainen et al., 2014). The ECG data for each subject was manually checked for physiological or computational artifacts like supraventricular extrasystoles or faulty peak detection, respectively. From RR interval time series (i.e., tachograms), we calculated the root mean square of successive differences (RMSSD) of adjacent RR intervals (Task Force of the European Society of Cardiology and the North American Society of Pacing Electrophysiology, 1996).

#### LEMON sample

Four minutes of resting ECG were acquired using a Biopac MP35 amplifier with the acquisition software AcqKnowledge version 4.0 (Biopac Systems Inc., http://www.biopac.com, Goleta, CA, USA) and three disposable electrodes on the thorax: the reference electrode was attached near the right collarbone, the measuring electrode on the left-hand side of the body on the same level as the 10^th^ rib, and the ground electrode on the right hip bone. The subjects were instructed to think about daily routines, relax, and breathe at a comfortable rate in sitting position. The peak detection and RMSSD calculation were performed using Kubios 2.2 (Tarvainen et al., 2014).

RMSSD values of our sample were natural log-transformed to obtain normally distributed data (Shapiro-Wilk tests; W = 0.99, p = 0.12). In the rest of the paper, log-transformed RMSSD will be referred to as “HRV”.

### 2.3. MRI acquisition

Brain imaging for both datasets was performed on the same 3T Siemens Magnetom Verio MR scanner (Siemens Medical Systems, Erlangen, Germany) with a standard 32-channel head coil. In both samples, subjects were instructed to keep their eyes open and not to fall asleep during the acquisition period.

#### LIFE sample

The structural T1-weighted images were acquired using a generalized auto-calibrating partially parallel acquisition technique (Griswold et al., 2002) and the Alzheimer’s Disease Neuroimaging Initiative standard protocol with the following parameters: inversion time (TI) = 900 ms, repetition time (TR) = 2.3 ms, echo time (TE) = 2.98 ms, flip angle (FA) = 9°, band width = 240 Hz/pixel, field of view (FOV) = 256 x 240 x 176 mm^3^, voxel size = 1 x 1 x 1 mm^3^, no interpolation. T2*-weighted functional images were acquired using an echo-planar-imaging (EPI) sequence with the following parameters: TR = 2000 ms, TE= 30 ms, FA = 90°, FOV = 192 x 192 x 144 mm^3^, voxel size = 3 mm x 3 mm, slice thickness = 4 mm, slice gap = 0.8 mm, 300 volumes, duration = 10.04 min. A gradient echo field map with the sample geometry was used for distortion correction (TR = 488 ms, TE 1 = 5.19 ms, TE 2 = 7.65 ms).

#### LEMON sample

The structural image was recorded using an MP2RAGE sequence (Marques et al., 2010) with the following parameters: TI 1 = 700 ms, TI 2 = 2500 ms, TR = 5000 ms, TE = 2.92 ms, FA 1 = 4°, FA 2 = 5°, band width = 240 Hz/pixel, FOV = 256 × 240 × 176 mm^3^, voxel size = 1 x 1 x 1 mm^3^. The functional images were acquired using a T2*-weighted multiband EPI sequence with the following parameters: TR = 1400 ms, TE = 30 ms, FA= 69°, FOV = 202 mm, voxel size = 2.3 x 2.3 x 2.3 mm^3^, slice thickness = 2.3 mm, slice gap = 0.67 mm, 657 volumes, multiband acceleration factor = 4, duration = 15.30 min. A gradient echo field map with the sample geometry was used for distortion correction (TR = 680 ms, TE 1 = 5.19 ms, TE 2 = 7.65 ms).

### 2.4. MR data preprocessing and analysis

#### Structural MRI

We analyzed structural brain alterations on the T1-weighted 3D image using VBM (Ashburner and Friston, 2000) as implemented in SPM12 (Wellcome Trust Centre for Neuroimaging, UCL, London, UK) and the Computational Anatomy Toolbox (CAT12: http://dbm.neuro.uni-jena.de/cat/), running on Matlab 9.3 (Mathworks, Natick, MA, USA). In the LEMON sample before the preprocessing, we removed the background noise from MP2RAGE on the computed uniform images via masking (Streitbürger et al., 2014). The preprocessing steps consisted of segmentation, bias-correction, and normalization using high-dimension Diffeomorphic Anatomical Registration Through Exponentiated Lie Algebra (DARTEL; Ashburner, 2007) with the template from 550 healthy controls of all ages in the IXI Dataset (http://www.brain-development.org) in MNI space. We then applied a 12-parameter affine registration and nonlinear transformation to correct for image size and position. The voxel size was resampled to 1.5 × 1.5 × 1.5 mm and smoothed using a 8-mm Gaussian kernel. For each subject, whole-brain gray matter volume (GMV) was calculated. An absolute threshold mask of 0.05 was specified in the analyses to cover the whole brain. For quality assessment, we visually inspected the segmentation quality and image homogeneity with the CAT12 toolbox. One participant from the middle-aged group was excluded because of MRI inhomogeneities.

#### Functional MRI

Preprocessing was implemented in Nipype (Gorgolewski et al., 2011), incorporating tools from FreeSurfer (Fischl, 2012), FSL (Jenkinson et al., 2012), AFNI (Cox, 1996), ANTs (Avants et al., 2011), CBS Tools (Bazin et al., 2014), and Nitime (Rokem et al., 2009). The pipeline comprised the following steps: (I) discarding the first five EPI volumes to allow for signal equilibration and steady state, (II) 3D motion correction (FSL mcflirt), (III) distortion correction (FSL fugue), (IV) rigid body co-registration of functional scans to the individual T1-weighted image (Freesurfer bbregister), (V) denoising including removal of 24 motion parameters (CPAC, Friston et al., 1996), motion, signal intensity spikes (Nipype rapidart), physiological noise in white matter and cerebrospinal fluid (CSF) (CompCor; Behzadi et al., 2007), together with linear and quadratic signal trends, (VI) band-pass filtering between 0.01-0.1 Hz (Nilearn), (VII) spatial normalization to MNI152 standard space (3 mm isotropic) via transformation parameters derived during structural preprocessing (ANTS). (VIII) The data were then spatially smoothed with a 6-mm FWHM Gaussian kernel.

The reproducible workflows containing all implementation details for our datasets can be found here: LIFE; https://github.com/fliem/LIFE_RS_preprocessing, LEMON; https://github.com/NeuroanatomyAndConnectivity/pipelines/releases/tag/v2.0

#### Eigenvector Centrality Mapping (ECM)

In ECM, each voxel in the brain receives a centrality value that is larger if the voxel is strongly correlated with many other voxels that are themselves central (Lohmann et al., 2010). ECM is computationally efficient, enables connectivity analysis at the voxel level, and does not require initial thresholding of connections (Lohmann et al., 2010). Here, the fast ECM implementation was used (Wink et al., 2012). We restricted our ECM analysis to GM, which we extracted with a mask from the tissue priors in SPM12 by selecting voxels with a GM tissue probability of 20% or higher. The resulting mask contained ~63,000 voxels covering the entire brain.

#### Exploratory Seed-based Functional Connectivity Analysis (SBCA)

To further explore the connectivity patterns of significant centrality changes across the whole brain, ECM was complemented by SBCA. Regions detected in ECM can be used as seeds in a subsequent SBCA to investigate intrinsic functional connectivity patterns (Taubert et al., 2011). A bilateral vmPFC seed was created by binarizing the significant ECM findings (MNI coordinates: [x=0, y=57, z=-6], cluster size k=62). Time series were extracted and averaged across all voxels of the seed. For each subject, a correlation between the time series of the seed and every other voxel in the brain was calculated using 3dfim+ (AFNI). The resulting correlation maps were Fisher r-to-z transformed using 3dcalc (AFNI).

#### Statistical analyses

Statistical analyses were carried out using the general linear model (GLM) approach implemented in SPM12. We performed one-way ANOVA with three age groups (young, middle, and old) as between-subjects factor together with HRV as the variable of interest and age, sex, study, and either total intracranial volume (TIV, for VBM analysis) or in-scanner head motion (mean FD; Power et al., 2012 for ECM and SBCA) as covariates of no interest.

We first calculated the interaction effect between HRV and age group. Based on the significant results of the ANOVA, we computed pairwise group differences using independent t-tests. Using one-sample t-tests, we further tested the main effect of HRV across all subjects, as well as for each age group separately. For each statistical analysis, a positive and a negative contrast were computed. Only results surviving whole-brain family-wise error (FWE) correction at p < 0.05 (cluster-level) with a voxel-level threshold of p < 0.001 were considered significant. All (unthresholded) statistical maps are available at NeuroVault (Gorgolewski et al., 2015) for detailed inspection in 3D (http://neurovault.org/collections/TELEUIIY).

### 2.5. Cognitive measurement and potential confounding factors for HRV

#### Sex

As HRV has been reported to differ between sexes (Koenig and Thayer, 2016; Voss et al., 2015), we analyzed sex differences in HRV per age group in a 2 (sex) × 3 (age group) ANOVA.

#### Smoking

Since smoking has a short- and long-term impact on HRV (Felber Dietrich et al., 2007; Hayano et al., 1990), we examined potential effects of smoking status on HRV. To this end, we classified subjects into three groups (smokers: N= 75, former smokers: N=84, and non-smokers: N=220, [no info available: NA=9]). We used a 2 (sex) x 3 (smoking) ANOVA to test the mean differences between the groups using sex as additional between-subjects factor.

#### Cognition

The Trail Making Test (TMT) is a cognitive test measuring executive function, including processing speed and mental flexibility. By drawing lines, subjects sequentially connect numbers and/or letters while their reaction times are recorded (Reitan, 1955; Reitan and Wolfson, 1995). In the first part of the test (TMT-A) the targets are all numbers (1, 2, 3, etc.), while in the second part (TMT-B), participants need to alternate between numbers and letters (1, A, 2, B, etc.). In both TMT A and B, the time to complete the task quantifies the performance and lower scores indicate better performance.

#### Blood Pressure

Systolic blood pressure (SBP) and diastolic blood pressure (DBP) were measured in a seated position using an automatic oscillometric blood pressure monitor (LIFE sample; OMRON 705IT, LEMON sample; OMRON M500) after a resting period of 5 min. While in the LIFE sample three consecutive blood pressure measurements were taken from the right arm in intervals of 3 minutes, in the LEMON sample measurements were taken from participants’ left arms on three separate occasions within two weeks. In each sample, all available measurements per participant were averaged to one systolic and one diastolic blood pressure value.

#### Anthropometric measurements

Subjects’ heights and weights were taken according to a standardized protocol by trained study staff. Body mass index (BMI; in kg/m^2^) was calculated by dividing the body weight by the square of the body height, while waist to hip ratio (WHR) was calculated as waist circumference measurement divided by hip circumference measurement (Huxley et al., 2010). As a control, all analyses on the association between HRV and the brain across the age groups were repeated with blood pressure (BP) and body mass index (BMI) as additional covariates of no interest.

For cognition, blood pressure, and anthropometric measurements, we assessed age-group differences statistically using one-way ANOVAs, and then tested their association with HRV using Spearman correlations for each age group. To determine statistical significance, we used a two-sided α-level of 0.05. Statistical analyses were conducted using R version 3.3.2 (R Core Team 2016).

## 3. Results

Details about the demographic, anthropometric, cardiovascular, and cognitive characteristics of the 388 participants can be found in Table 1. The age groups differed significantly in all variables (Table 1).

There was a significant main effect of age group on HRV (*F*(2,382) = 63.552, p =2×10^−16^, η^2^ = 0.182), sex did not show a significant main effect on HRV (*F*(1,382) = 0.187, p = 0.666), and there was no significant age group × sex interaction on HRV (*F*(2,382) = 0.233, p = 0.792). HRVs of smokers, former smokers, and non-smokers did not differ significantly from each other (main effect smoking group: *F*(2,373) = 1.241, p = 0.290, main effect of sex: *F*(1,373) = 0.473, p = 0.492; smoking group × sex interaction: *F*(2,373) = 0.606, p = 0.546).

HRV was negatively correlated with age (rho = −0.21, p = 0.010), BMI (rho = −0.207, p = 0.020), and DBP (rho = −0.231, p = 0.012) within the middle-aged individuals. No significant associations were found between HRV and mean FD, SBP, WHR, TMT A, and TMT B in any of the age groups (Supplementary Table 1).

### Voxel-based Morphometry (VBM)

There was no significant association between HRV and GMV across all subjects. Also, an ANOVA did not yield a significant age group x HRV interaction on GMV. While an exploratory one-sample t-test in the middle-aged group indicated a significant HRV-related increase of GMV in left cerebellum (MNI coordinates: [−15, −87, −51], k = 1540, T = 3.92, pFWE = 0.004), there were no significant effects of HRV on GMV for younger and older adults. Control analyses that included BP and BMI as covariates of no interest did not change the results.

### Eigenvector Centrality Mapping (ECM)

A significant effect of age group on the relation between resting HRV and EC was detected in the bilateral vmPFC (MNI coordinates: [0, 57, −6], k = 62, F = 10.79). The beta values from bilateral vmPFC for each age group are plotted in Figure 1A, suggesting that younger adults show a stronger association between HRV and EC in bilateral vmPFC than middle-aged and older individuals (Table 2). This was supported by post-hoc two-sample t-tests, which indicated that the correlation between HRV and EC in bilateral vmPFC was significantly stronger for the contrasts of young > old and young > middle-age (Table 2). A one-sample t-test across all subjects showed increased EC with higher HRV in bilateral PCC (Figure 1B). The negative contrast did not yield any significant results. In separate one-sample t-tests for each age group, we found HRV-dependent EC increases in right vmPFC, bilateral PCC, and superior frontal gyrus (SFG), as well as HRV-dependent EC decreases in left superior occipital gyrus (SOG) including cuneus and calcarine sulcus in the group of young subjects. Our data did not show any significant positive or negative correlation with HRV in the groups of middle-aged and old subjects that were correctable for multiple comparisons. The complete ECM results are presented in Table 2. Control analyses that included BP and BMI as covariates of no interest did not change the results.

**Fig. 1.**
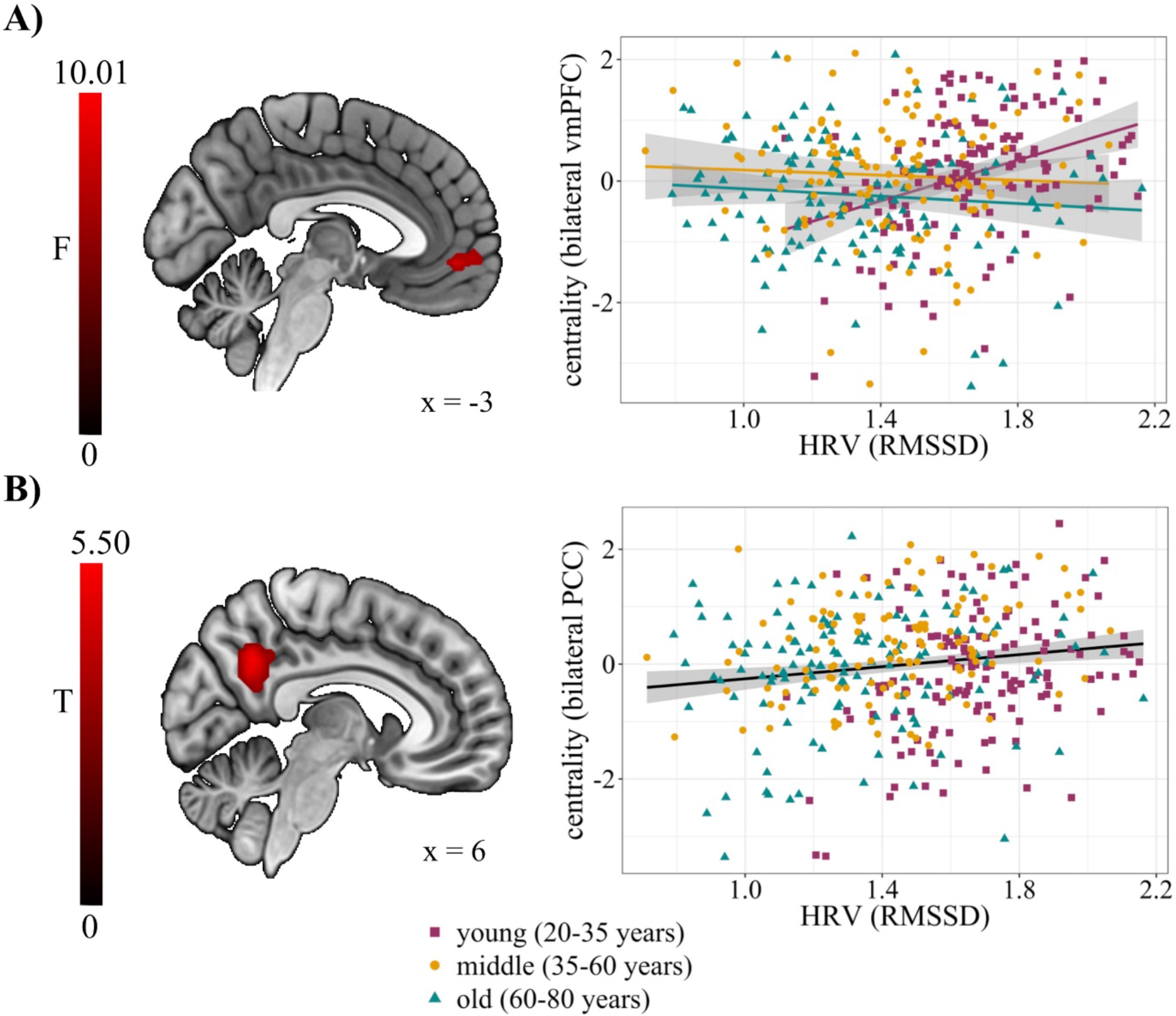
Association between resting heart rate variability (HRV), measured as root mean square of successive differences (RMSSD), and eigenvector centrality (EC). **A)** The interaction between age group and HRV was significant in the bilateral ventromedial prefrontal cortex (vmPFC; MNI coordinates: [0, 57, −6], k = 62, F = 10.79, pFWE = 0.006), displayed at x = −3. **B)** An increased EC in the bilateral posterior cingulate cortex (PCC; MNI coordinates [6, −54, 36], k = 204, T = 5.29, pFWE < 0.001) across all age groups, displayed at x = 6. Results are shown at a voxel threshold of p < 0.001 with family-wise error (FWE) correction with p < 0.05 at the cluster level.

**Table 2.**
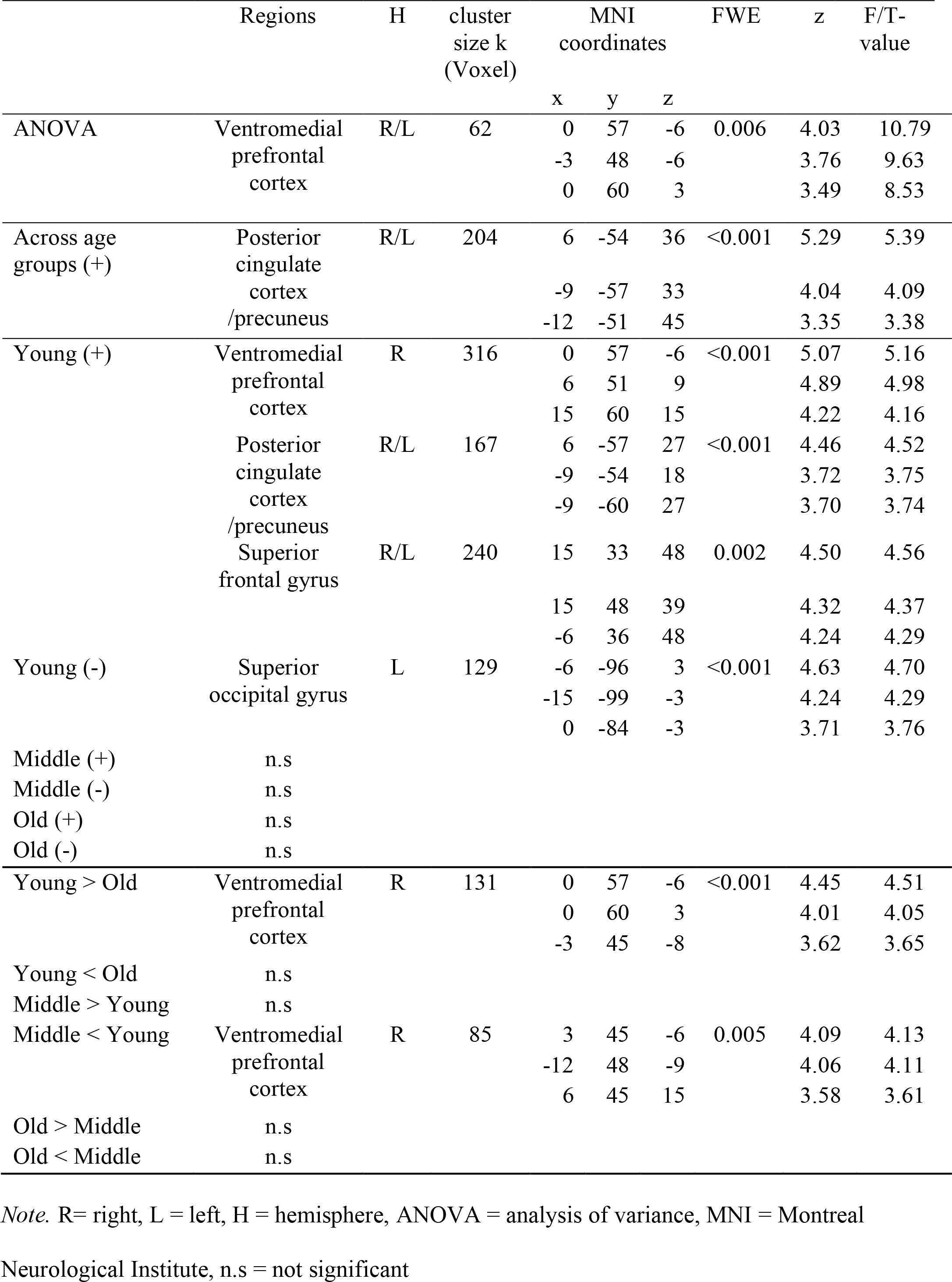
Brain regions that show significant increases or decreases in eigenvector centrality with heart rate variability (HRV). Thresholds: p < 0.001 at the voxel and p < 0.05 with family-wise error (FWE) correction at the cluster level.

### Exploratory Seed-based Functional Connectivity Analysis (SBCA)

In the additional exploratory seed-based functional connectivity analysis, a significant effect of age group on the relation between resting HRV and whole-brain bilateral vmPFC connectivity was found in bilateral cerebellum, right superior parietal lobe (SPL), left middle occipital gyrus (MOG) and inferior occipital gyrus (IOG), and left SFG extended to supplementary motor area (SMA). The beta values from the right cerebellum for each age group are plotted in Figure 2A, suggesting that younger adults show stronger functional connectivity between bilateral vmPFC and right cerebellum than middle-aged and older individuals (Table 3). The post-hoc two-sample t-tests similarly indicated that higher HRV levels were significantly correlated with stronger functional connectivity between bilateral vmPFC and bilateral cerebellum, right SPL, left MOG, left post-central gyrus, and left SMA for the contrasts of young > old and young > middle (Table 3). A one-sample t-test in the overall sample, to assess the association between HRV and bilateral vmPFC connectivity, showed an increased functional connectivity with left middle frontal gyrus (MFG) extending to dorsolateral prefrontal cortex (DLPFC) (Figure 2B). Separate one-sample t-tests for each age group showed no significant association for the middle-aged and old subjects but an increased vmPFC connectivity in distributed brain regions including bilateral cerebellum, bilateral MOG, and right SMA for the young subjects. We did not observe any significant negative correlations neither in the overall sample nor in each age group. Control analyses that included BP and BMI as covariates of no interest did not change the results. The complete SBCA results are presented in Table 3.

**Fig. 2.**
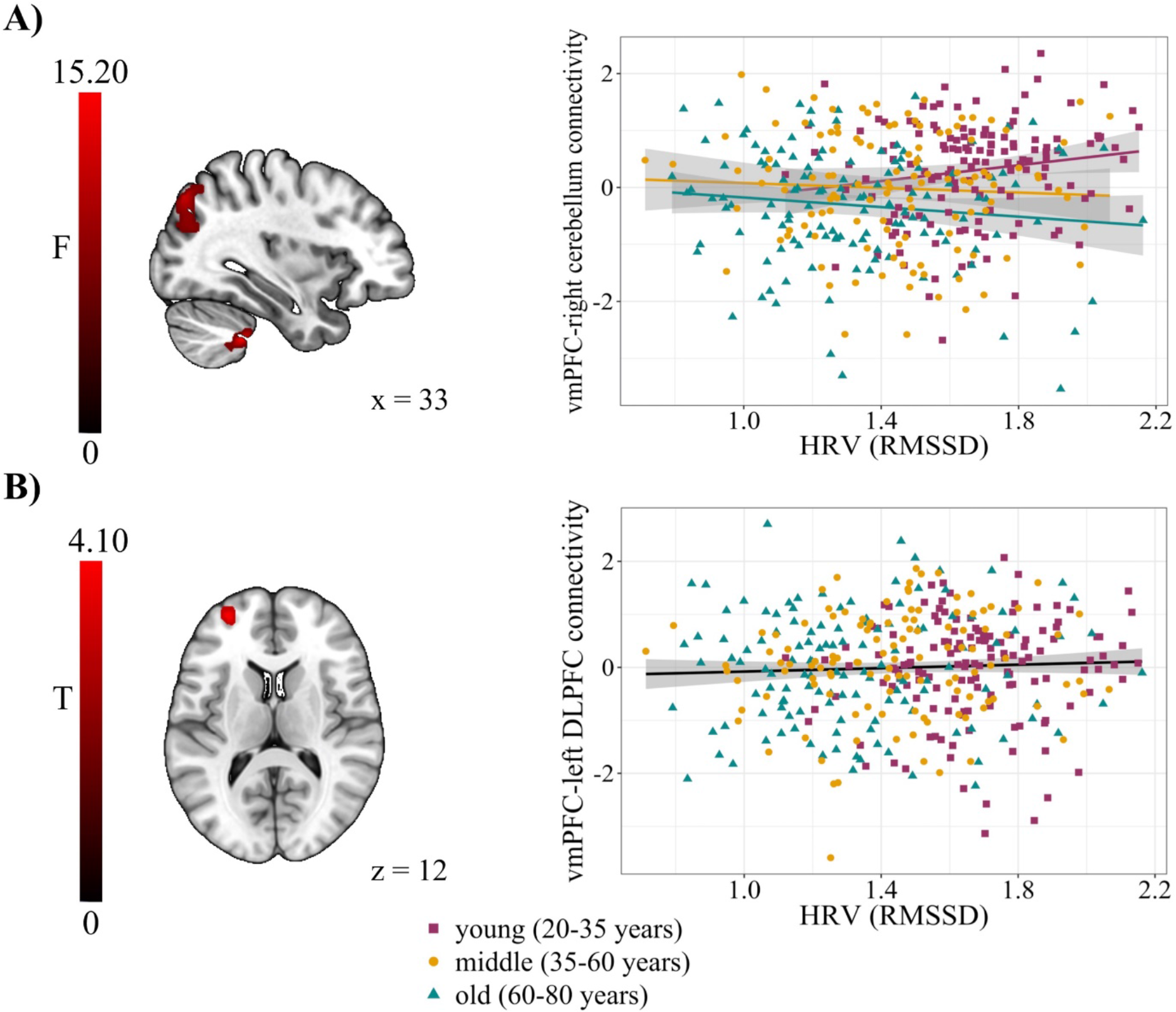
Association between resting heart rate variability (HRV), measured as root mean square of successive differences (RMSSD), and brain function in an exploratory seed-based functional connectivity analysis originating from bilateral ventromedial prefrontal cortex (vmPFC). **A)** The interaction between age group and HRV was significant in the right cerebellum (MNI coordinates [33, −42, −45], k = 46, F = 15.19, pFWE < 0.001), displayed at x = 33. **B)** An increased functional connectivity in the right dorsolateral prefrontal cortex (DLPFC; MNI coordinates [-30, 54, 12], k = 67, T = 4.10, pFWE = 0.032) was found across all age groups, displayed at z = 12. Results are shown at a voxel threshold of p < 0.001 with family-wise error (FWE) correction with p < 0.05 at the cluster level.

**Table 3.**
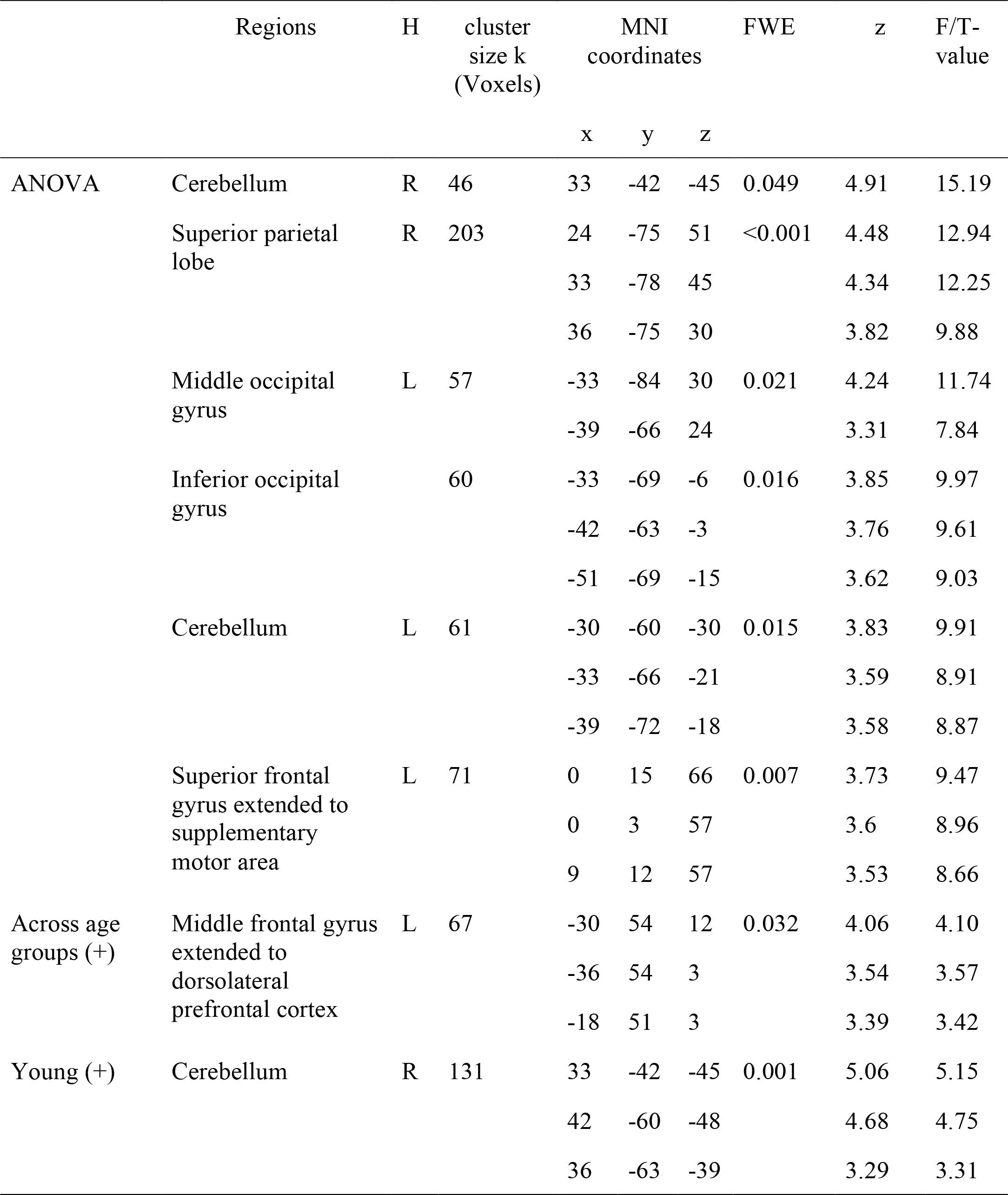
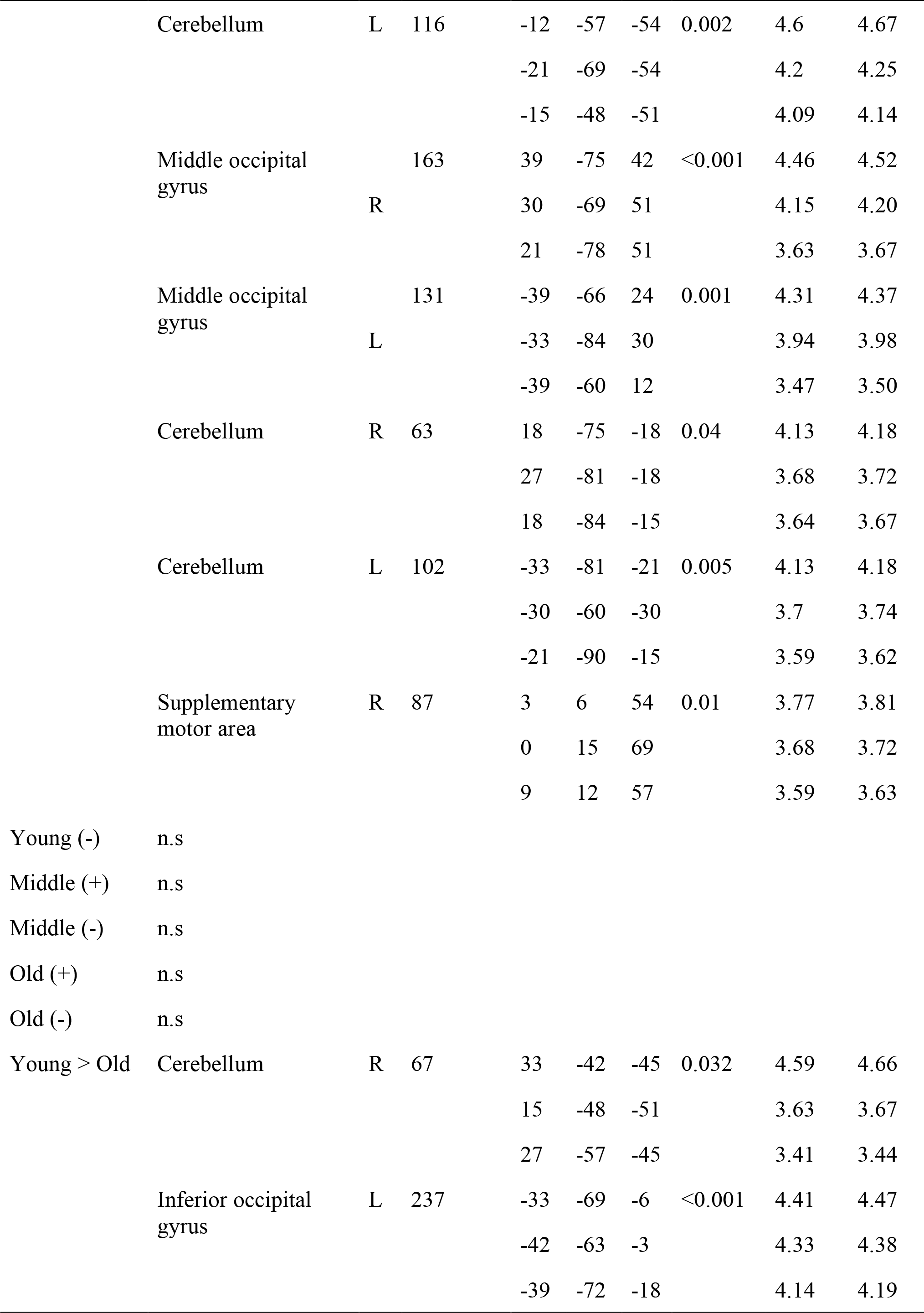
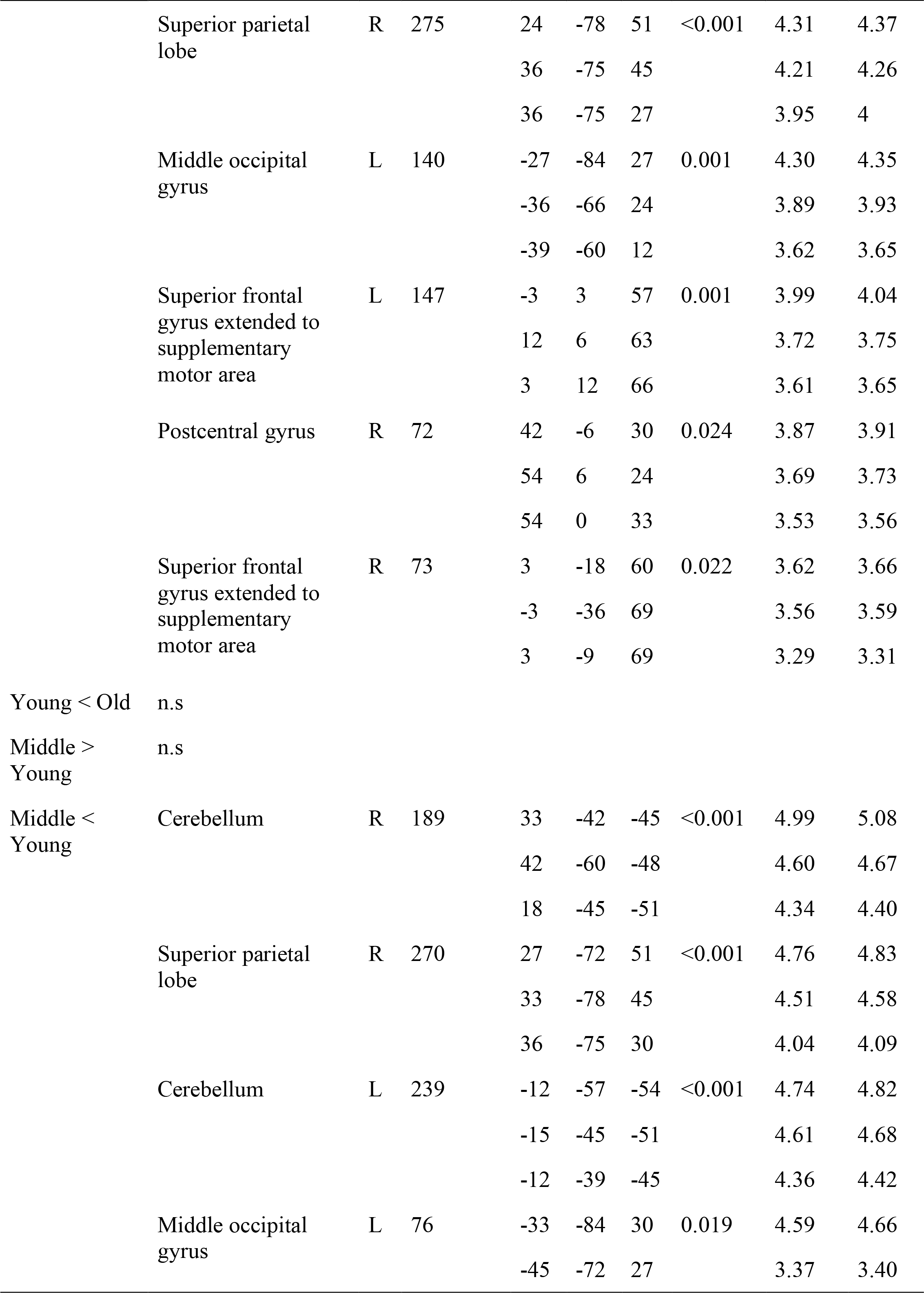
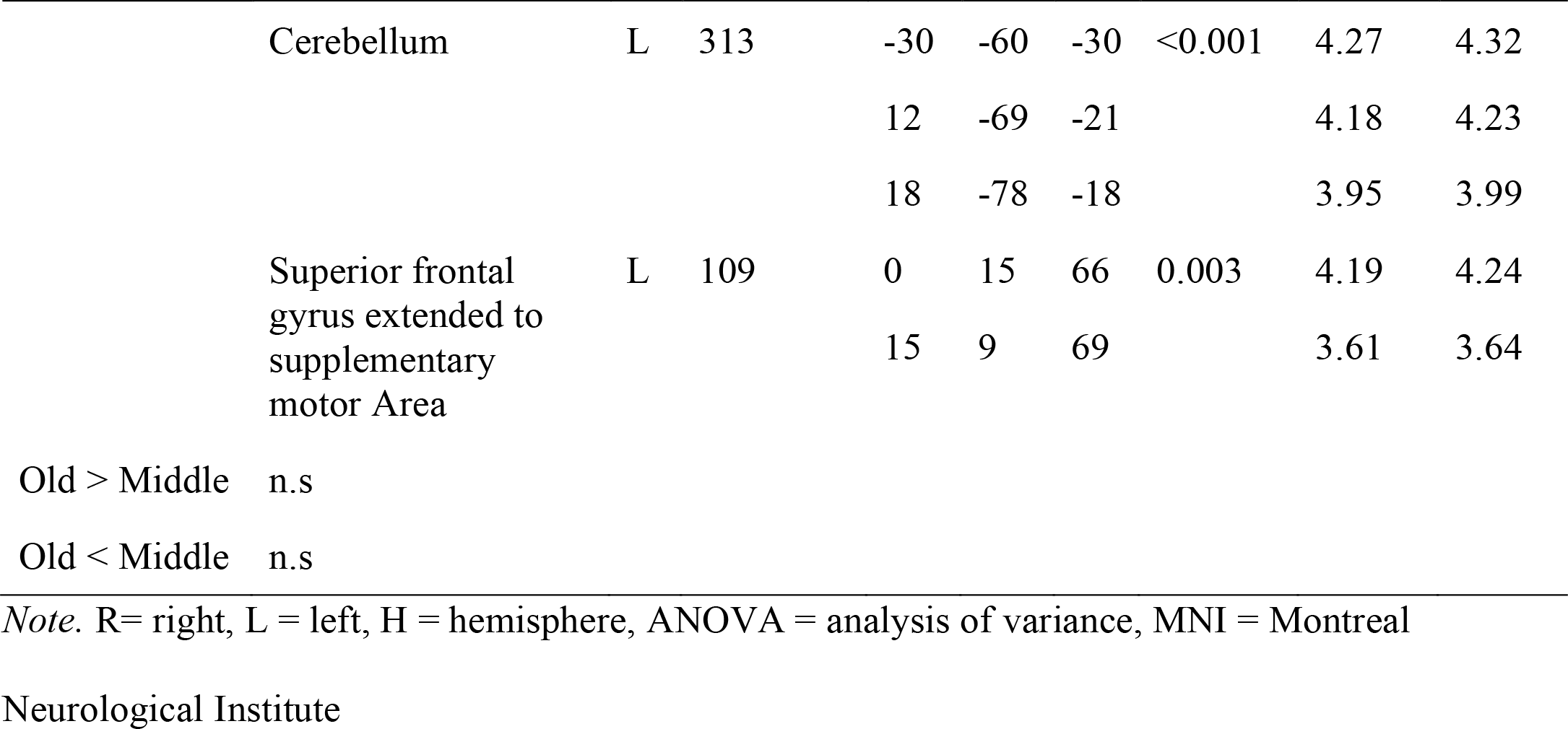
Brain regions that show resting heart rate variability-related connectivity with the bilateral ventromedial prefrontal cortex (vmPFC) in an exploratory seed-based functional connectivity analysis. Thresholds: p < 0.001 at the voxel and p < 0.05 with family-wise error (FWE) correction at the cluster level.

## 4. Discussion

In the present study, we assessed the relationship between parasympathetic cardioregulation (using resting HRV) and brain structure (using VBM) as well as whole-brain resting-state connectivity (using ECM and SBCA) in a large sample of healthy young, middle-aged, and old participants. We found the frequently observed age-related decrease in resting HRV (Almeida-Santos et al., 2016; Voss et al., 2015) to be accompanied by alterations in brain function. Specifically, higher HRV was linked to stronger network centrality in several brain regions, particularly along the cortical midline. In the PCC, this correlation was present in all age groups while in the vmPFC, network centrality was related to higher HRV in young but not in middle-aged and old adults. These findings support the view that altered HRV during aging may have a functional brain component associated with it.

### 4.1. Age-dependent association of resting HRV with functional connectivity

Given the relationship between HRV and age (Almeida-Santos et al., 2016; Voss et al., 2015), HRV and brain structure (Wei et al., 2018), as well as HRV and brain function (Sakaki et al., 2016), we hypothesized the neural correlates of resting HRV to be also age-dependent. Our results confirm that the relationship between HRV and network centrality at rest differs between age groups. Evidence is accumulating that alterations of intrinsic brain activity are a key feature of normal brain aging (Damoiseaux et al., 2008): age-dependent intrinsic connectivity alterations in the DMN have been found not only in healthy aging (Ferreira and Busatto, 2013) but also in (age-related) pathologies, for example, in individuals with a high familial risk for depression (Posner et al., 2016) and in young APOE-ε4 carriers (Filippini et al., 2009), which is a possible biomarker for Alzheimer’s dementia (Kanekiyo et al., 2014). Our results that resting HRV is related to increased network centrality in medial frontal regions in the young but not in the middle-aged and old age group could be interpreted in the framework of the functional plasticity hypothesis of cognitive aging (Greenwood, 2007). According to this hypothesis, the structural vulnerability particularly of prefrontal cortex leads to an age-related functional reorganization (e.g., Grady, 2012; for a detailed review). Changes in the resting-state network architecture around the vmPFC that are related to parasympathetic cardioregulation could thus represent altered cardiovascular control with advancing age and concomitant network reorganization.

In addition to the age-dependent association of resting HRV with functional brain network centrality in medial frontal regions, we also found an HRV-related bilateral medial parietal cluster in the PCC that was independent of age. Both vmPFC and PCC are central nodes of the DMN (Greicius et al., 2003; Uddin et al., 2009) and have been related to self-generated or internally directed mental processes like thoughts and feelings (Andrews-Hanna et al., 2014; Raichle et al., 2001). Regions of the DMN – and particularly its medial frontal (e.g., vmPFC) and parietal components (e.g., PCC and precuneus) – have also been implied in the central processing of autonomic – and particularly parasympathetic – function (Beissner et al., 2013; Benarroch, 1993). It is plausible that in the absence of external stimulation, brain function (i.e., activity and connectivity) is predominantly allocated to the “internal milieu”, that is, to monitoring and regulating bodily signals (e.g., the parasympathetic “rest-and-digest”). Fittingly, the PCC has been found active in tasks that involved the assessment of self-relevance (Yu et al., 2011) as well as self-location and body ownership (Guterstam et al., 2015), while the vmPFC was related to processing bodily information (Gusnard et al., 2001), autonomic control (Critchley et al., 2011), and cardiovascular arousal (Wong et al., 2007). Similarly, a causal role of the PFC for cardiovascular activity (e.g., HR and HRV) was found in a meta-analysis of non-invasive brain stimulation and autonomic functioning (Makovac et al., 2017).

The exploratory SBCA similarly showed an age-dependent relationship between resting HRV and functional brain connectivity. Specifically, we found stronger functional connectivity between bilateral vmPFC and a widespread set of brain regions including bilateral cerebellum, occipital gyrus, right SPL, and SFG extending to SMA in young but not middle-aged and old adults. These results extend the ECM findings by suggesting additional cortico-cerebellar regions might be involved in the modulation of visceral processes. In line with this interpretation, activation in the cerebellum has been connected to the regulation of visceral responses (Demirtas-Tatlidede et al., 2011), fear conditioning (Leaton, 2003; Sacchetti et al., 2002), feeding (Tataranni et al., 1999), as well as the coordination and control of cardiovascular activities (Bradley et al., 1991; Ghelarducci and Sebastiani, 1996). Furthermore, autonomic activity during cognitive and motor tasks was positively associated with activation in the cerebellum and, among other regions, the SMA and dorsal ACC (Critchley et al., 2003).

Despite previous evidence of the relationship between brain structure and vagally-mediated HRV in central autonomic network regions (Wei et al., 2018), using whole-brain VBM analysis, we only found a significant GMV change related with resting HRV in the cerebellum for the middle-aged group. Notably, in the study by Wei et al. (2018) reduced GM volume in the cerebellum was associated with HR (but not HRV) in healthy middle-aged individuals. The divergent results could be due to different measurement parameters (e.g., MRI sequence parameters) but also to different effect size and statistical power (for more details see *Limitations*).

### 4.2. Physiological and psychophysiological interpretations of HRV

The most fundamental (purely physiological) understanding of the role of the ANS – and particularly the PNS – is to ensure visceral and cardiovascular functioning or bodily homeostasis – by allowing rapid adaptive behavioral and physiological reactions in ever-changing environments or by disengagement and relaxation in resting moments (“rest-and-digest”; e.g., Cannon, 1929).

More *psycho*physiological interpretations of ANS function have extended this view to cognitive, affective, and social phenomena: For example, the two main theoretical perspectives of HRV – the polyvagal theory (Porges, 2007, 2001, 1995) and the neurovisceral integration model (Smith et al., 2017; Thayer and Lane, 2000; Thayer and Ruiz-Padial, 2006) – suggest that PNS activation can serve as a biomarker of what can be summarized as “top-down” self-regulation (Holzman and Bridgett, 2017). The polyvagal theory (Porges, 2007) takes an evolutionary approach, according to which the role of the ANS and particularly the vagus nerve can be understood as increasing adaptability through socially engaged behaviors (e.g., self-soothing) and inhibition of sympathetic-adrenal influences on the body (Porges, 2007). The neurovisceral integration model (Smith et al., 2017; Thayer and Lane, 2000; Thayer and Ruiz-Padial, 2006) highlights the role of vagally-mediated HRV for emotional or self-regulation. This model explicitly links the brain and the rest of the body by assuming that the PFC – and particularly the vmPFC – tonically inhibits the amygdala, which affects autonomic function, thereby linking both nervous systems to inhibitory or (self-)regulatory processes (Kemp et al., 2017; Thayer et al., 2012). Convergently, resting HRV has recently been associated with vmPFC activation during a dietary self-control task in young adults (Maier and Hare, 2017). Both theories draw on evidence that higher HRV is indicative of better bodily functioning by enabling physiological and behavioral adaptation through cognitive and socio-emotional flexibility. Taken together, our findings are consistent with both the polyvagal theory and the neurovisceral integration model of HRV and extend them by providing evidence for a brain network component of vagally-mediated HRV in healthy aging.

## 5. Limitations

There are a number of limitations that should be considered in the interpretation of our results. The study design is cross-sectional and does not allow us to infer the directionality of the association between resting HRV and the brain. Additionally, our health criteria also allowed inclusion of subjects with higher BMI (>25 kg/m^2^) or untreated/undiagnosed hypertension (SBP > 140 mmHg, DBP > 90 mmHg). This makes it difficult to disentangle HRV-related influences from other bodily/cardiovascular influences – which are also physiologically related (BMI: Molfino et al., 2009; BP: Singh et al., 1998). However, control analyses that accounted for BP and BMI showed very similar results of the association between resting HRV and the brain. Although psychological interpretations of a single physiological marker like resting HRV are intrinsically limited, previous studies have associated HRV with different trait or state levels of, for example, executive control (Capuana et al., 2014), stress (Sin et al., 2016), and emotion regulation (Williams et al., 2015). For a psychological interpretation of our finding that the association between HRV and functional connectivity at rest is age-dependent, similar analyses on task-related parasympathetic and neural activity could be helpful. Although we accounted for systematic study differences in the second-level GLM, different acquisition parameters of the rs-f/MRI and ECG may have influenced our results (e.g., for structural MRI; Streitbürger et al., 2014). Further, we calculated the RMSSD using 10 s of ECG data, which has been shown to be a valid measurement (Munoz et al., 2015; Nussinovitch et al., 2011a, 2011b). Nevertheless, ECG data recorded over longer periods (e.g., 24-hour) can complement this “ultra-short” evaluation of parasympathetic function.

## 6. Conclusion

In this cross-sectional study, we examined the association of resting HRV with brain structure and functional brain connectivity in different age groups of healthy adults. Our main findings are correlations between resting HRV and brain network architecture in the PCC across all age groups and in the vmPFC in young but not in middle-aged and old subjects.

These support the view that the well-known HRV decrease with age may have a functional brain network component along the cortical midline. Consistent with the role of these areas in affective, cognitive, and autonomic regulation, our results provide a comprehensive picture of the differential effect of age on heart-brain interactions and extend our knowledge of parasympathetic cardioregulation being important for healthy aging.

## 8. Acknowledgments

This study was supported by LIFE Leipzig Research Center for Civilization Diseases at the University of Leipzig funded by the European Union, European Regional Development Fund, and the Free State of Saxony. The authors would like to thank all volunteers for their participation in one of the two studies. Further, we thank all researchers, technicians and students who planned, collected, entered and curated data, used in this manuscript.

---

**Supplementary Table 1.** Association (Spearman’s rho) between heart rate variability (HRV) and other parameters per age group.

|  | Young | Middle | Old |
| --- | --- | --- | --- |
| Age | -0.10 | -0.21* | -0.01 |
| mean FD (mm) | -0.13 | -0.11 | 0.09 |
| BMI (kg/m²) | 0.13 | -0.20* | -0.14 |
| WHR | -0.04 | -0.17 | -0.08 |
| SBP (mmHg) | 0.01 | -0.17 | -0.01 |
| DBP (mmHg) | -0.06 | -0.23* | -0.01 |
| TMT A (s) | 0.01 | -0.12 | -0.04 |
| TMT B (s) | -0.06 | -0.12 | 0.02 |

*\*p < 0.05; \*\*p < 0.01; \*\*\*p < 0.001, 2-tailed.*
*Note.* FD = framewise displacement; BMI = body mass index; WHR = waist to hip ratio; SBP
= systolic blood pressure; DBP = diastolic blood pressure, TMT = trail making test.

